# Neuromark PET: A multivariate method for estimating whole brain fMRI guided intrinsic networks and connectomes from fMRI and PET data

**DOI:** 10.1101/2024.01.10.575131

**Authors:** Debbrata K. Saha, Anastasia Bohsali, Rekha Saha, Ihab Hajjar, Vince D. Calhoun

## Abstract

Positron emission tomography (PET) and magnetic resonance imaging (MRI) are both widely used neuroimaging techniques to study brain functional and molecular connectivity. Although whole brain resting functional MRI (fMRI) connectomes (a matrix describing the inter-regional connectivity patterns) are widely used, the integration or association of whole brain molecular connectomes with PET data are rarely done. This likely stems from the fact that PET data is typically analyzed by using a region of interest approach, while whole brain spatial networks and their connectivity (covariation) receive much less attention. As a result, to date, there have been little focus on directly comparing whole brain PET and fMRI connectomes. In this study, we present a method that uses spatially constrained independent component analysis (scICA) (utilizing fMRI components as spatial priors) to estimate corresponding (Amyloid) PET and fMRI connectomes and examine the relationship between them using datasets that include individuals with mild cognitive impairment (MCI). Our results demonstrate highly modularized PET connectome patterns that complement those identified from resting fMRI. In particular, fMRI showed strong intra-domain connectivity with inter-domain anticorrelation in sensorimotor and visual domains as well as default mode network. PET amyloid data showed similar strong intra-domain effects, but showed much higher correlations within cognitive control and default mode domains, as well as anticorrelation between cerebellum and other domains. The estimated fMRI informed PET networks have similar, but not identical, network spatial patterns to the resting fMRI networks, with the fMRI informed PET networks being slightly smoother and, in some cases, showing variations in subnodes. To further compare the two modalities, we also analyzed the differences between individuals with MCI receiving medication versus a placebo. Results show both common and modality specific treatment effects on fMRI and PET connectomes. From our fMRI analysis, we observed higher connectivity differences in various regions, such as the connection between the thalamus and middle occipital gyrus, as well as the insula and right middle occipital gyrus. Meanwhile, the PET analysis revealed increased activation between the anterior cingulate cortex and the left inferior parietal lobe, along with other regions, in individuals who received medication versus placebo. In sum, our novel approach identifies corresponding whole-brain fMRI informed PET and fMRI networks and connectomes. While we observed common patterns of network connectivity, our analysis of the MCI treatment and placebo groups revealed that each modality captures modality and group specific information about brain networks, highlighting differences between the two groups in both network expression and network connectivity.

## 1. Introduction

Brain network connectivity (functional or molecular) which describes inter-network coupling or covariation, is a widely utilized tool for inferring connections among various brain networks. While certain imaging techniques are restricted to analyzing a limited number of regions of interest, comprehensively examining spatial networks and their connectivity across the entire brain can be studied via the whole brain connectome [1].

To investigate how brain function supports mental tasks, cognitive neuroscience integrates the experimental methods of cognitive psychology with a number of different imaging techniques. One of the commonly used imaging techniques in cognitive neuroscience is fMRI, which measures the changes in blood oxygenation levels associated with neural activity [2, 3]. fMRI has been successfully used to extract whole brain functional connectivity (FC) among brain regions. A related concept, functional network connectivity (FNC) is a measure of the temporal covariation among functional connectivity networks obtained from fMRI time series data (e.g. intrinsic networks estimated via independent component analysis). PET is another widely used brain molecular imaging technique that provides unique insight into the brain by measuring specific tracer information, such as amyloid uptake. PET can be analyzed quantitatively to study the tracer uptake in regions or volumes. An alternative approach, which is receiving renewed interest, is to analyze molecular network connectivity (MNC), which maps statistical dependencies (or coupling) between brain networks based on molecular activity, revealing pathways of disease propagation and functional interactions [4, 5, 6]. PET analyses predominantly focuses on analyzing regional or global summaries of brain activity [7, 8, 9]. Network based analysis is typically confined to inter-subject covariation among a few regions of interest. Consequently, research on estimating whole-brain spatial networks and the connectivity between them through PET data remains scarce.

There is growing research interest in analyzing patients with mild cognitive impairment (MCI) using various imaging techniques. MCI may indicate an initial stage of Alzheimer’s disease (AD) and is characterized by cognitive dysfunction in individuals who have not yet reached the dementia threshold [10, 11, 12]. Individuals with MCI are at an elevated risk of developing dementia, particularly AD. Research conducted in specialized medical facilities has indicated that MCI patients progress to AD at an annual rate of 10% to 15%, and roughly 80% of these patients develop AD within about six years of being under observation [13, 14]. As the conversion rate from mild cognitive impairment to AD is significantly higher than that of healthy age-matched individuals, early identification of individuals at risk for Alzheimer’s disease is crucial. Such identification would facilitate the targeted application of available treatments to the appropriate group.

A number of studies have reported the efficacy of PET and fMRI data in analyzing patients with MCI/Alzheimer’s disease. fMRI studies have found aberrant functional connectivity in default mode network (DMN), sensorimotor network (SMN), and dorsal attention (DAN) in AD patients compared to healthy individuals [15]. Additionally, extensive analysis of MCI patients has reported the loss of DMN integrity [16, 17], decreased correlations in temporal regions (hippocampus, thalamus, fusiform gyrus), and the prefrontal cortex (inferior frontal gyrus, orbital, olfactory cortex, superior frontal gyrus, medial, middle frontal gyrus) [18]. Recent fMRI studies have consistently reported reduced functional connectivity in the DMN, salience network, dorsal attention network, and frontoparietal network for both MCI and AD patients [16, 19, 20, 21]. PET has also been successfully employed to distinguish subjects with AD from healthy controls [22, 23], analyze patients with MCI [24, 25, 26], as well as to differentiate other forms of dementia, such as frontotemporal dementia (FTD) [27], and mental disorders like SZ [28]. PET has proven itself to be an extremely useful neuroimaging tool to investigate neural network activation and connectivity patterns. However, to our knowledge, no studies have thus far examined covariation between brain networks identified by PET by utilizing scICA with prior brain template, nor have there been studies directly comparing PET and fMRI whole brain connectomes in either neurologically normal participants or patient populations. In this work, we present a novel method for estimating and comparing brain connectomes using PET and fMRI data.

Our approach involves applying scICA with fMRI components as a prior brain template to both modalities, resulting in corresponding modality specific spatial networks for each. We then compute the FNC (i.e., the connectome) by calculating temporal cross correlation for the fMRI data, while for the PET data, we compute MNC via inter-subject covariation. The connectivity patterns for each modality are computed separately, and then compared to evaluate similarities and differences between FNC and MNC; and the spatial networks (connectivity networks). To evaluate the similarities and differences in the FNC and MNC effects, we also analyzed differences between individuals having both a PET and fMRI scan and with MCI treated with either candesartan or a placebo. To the best of our knowledge, our proposed method is the first approach to leverage a constrained data-driven framework to compare whole brain connectomes in PET versus fMRI, and in particular our approach provides a powerful framework that can be extended in future work.

The structure of this research article is as follows: The first section, Material and Methods, provides details on data acquisition and post-processing. This is followed by a detailed explanation of the proposed scICA method, which includes information on pairwise correlation and group activation procedures used in the study. The Experimental Results section presents an extensive validation of the performance of the proposed scICA, using a comparative analysis of results from PET and fMRI analyses. Methodological issues and findings are discussed in the Discussion section. Finally, we offer closing thoughts and an overall summary of the work in the Conclusion section.

## 2. Materials and Methods

### 2.1. Data Acquisition and Preprocessing

This study included MRI and Amyloid PET brain imaging obtained at baseline of the double-blind randomized controlled clinical trial, CEDAR (Candesartan’s Effects on Alzheimer’s Disease and Related Biomarkers; ClinicalTrials.gov Identifier: NCT02646982). The trial recruited non-hypertensive participants 50 years or older (mean age: 68.1 years; n=48, 62% women; n=15, 20% African American) with MCI using Petersen et al criteria [29] and evidence of AD pathology measured by CSF *Aβ* and tau or amyloid PET (prodromal AD). Participants were stratified by current use of symptomatic treatment and randomized to escalating doses of Candesartan or placebo and treated for 1 year and both resting fMRI and PET data were collected. This study collected data at the baseline and after 12 months. In this methods feasibility study, to maximize the data, we considered both the baseline and 12 month scans. Overall FNC/MNC and network spatial map comparisons between fMRI and PET were computed at only the baseline scans.

#### 2.1.1. PET data

In this study, we used 11*C − PiB* imaging which provides an estimate of the accumulation of *Aβ* throughout the brain. Radiotracers were prepared by the Emory radiopharmacy, and imaging was done at the Emory Center for Systems Imaging Core (CSIC) using a High-Resolution Research Tomography (HRRT) (CTI, Inc. Knoxville, TN). PET images were taken for a duration of 20 minutes, which consisted of 4 *×* 5 minutes frames. These images were obtained approximately 50 minutes after intra-venous administration of 14.81 +/-0.88 mCi of 11C-PiB or approximately 80 minutes after intravenous administration of 9.72 +/-0.69 mCi of 18F flortaucipir. The images were processed using several steps. First, they were smoothed using a Gaussian kernel with a full width at half max of 10 mm. Next, they were resampled to a 2 *×* 2 *×* 2 mm resolution and then averaged over frames using the AFNI software package ^1^ [30]. Preprocessed PET data were subjected to additional analysis using the PETSurfer pro-cessing pipeline. To be more precise, each subject’s PET and T1-weighted scans were co-registered using PETSurfer and a nonlinear warp. After that, Z-score normalization was applied to the PET data before running scICA.

#### 2.1.2. fMRI data

The fMRI acquisition used a Siemens Magnetom Prisma Fit 3T scanner to collect resting-state functional MRI (rs-fMRI) data. The imaging volumes were acquired with 48 transversal slices with a field of view of 220 mm (AP) *−* 144 mm(FH) *−* 220 mm (RL), and a voxel resolution of 3 *×* 3 *×* 3 mm, and other parameters like FA=90^◦^, TE=27ms, TR=2500ms. BrainForge, a cloud-enabled, web-based analysis platform for neuroimaging research, was used to process the rs-fMRI data [31]. The data analysis was done using a standardized SPM12-based analysis pipeline, which included several steps as slice timing correction, head-motion correction, spatial normalization to a tissue probability map (TPM) MNI template, and smoothing using a 6mm FWHM Gaussian kernel [32].

### 2.2. Spatially Constrained ICA

ICA has been extensively utilized in biomedical domains to analyze the brain activity [33, 34, 35, 36, 37, 38, 39]. In fMRI data, ICA is typically applied on *N × M* data matrix *X*, where *N* and *M* represent the time points and voxels respectively. From the ICA decomposition, we get the *N × N* linear mixing matrix and *N × M* independent component matrix. In structural MRI (or PET) data, ICA is applied to the subject volume matrix *X*, and decompose into a mixing matrix *A*, and a source matrix *S*. The mixing matrix represents the relationship between subjects and sources, and source matrix preserves the relationship between sources and voxels of brain. The ICA decomposition effectively performs spatial filtering to handle noise and provides maximally spatially independent sources, each of which exhibit inter-subject covariation [40, 41, 42]. The loading parameters provide information about the degree to which each component is expressed in a given subject. A number of prior studies have applied ICA to PET data [43, 44, 45] including some that compare networks for fMRI and PET data, or utilize data fusion approaches such as joint ICA to extract jointly covarying networks. However these studies do not typically compare the resulting connectomes of fMRI and PET, and they typically work with lower dimensions of components (e.g., 20) rather than a higher dimensional space (which helps us to visualize the modularity among the networks). In this work we leverage an existing set of 53 resting networks from a 100 component decomposition as spatial priors using spatially constrained ICA, and compare the spatial maps and connectome results for PET and fMRI.

Constrained ICA is an extension of ICA that isolates targeted sources from the decomposition process by using prior knowledge. First, a reference *R* is chosen for the prior information that includes the desired sources. Then, a closeness measure *g_W_* (*S, R*) between an estimated signal *S* = *WX* and the reference signal *R* is defined to restrict the learning process [46, 47]. The spatially constrained ICA (scICA) framework is formulated via the following augmented multi-objective loss [48]:

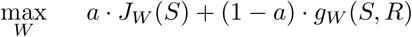

where, *J_W_*(*S*) denotes a normalized approximation of the negentropy of the estimated source *S* (this is a standard contrast function for ICA), *g_W_* (*S, R*) = 𝔼(*S, R*) is the Pearson correlation coefficient between *S* and *R*, and *a* is a hyperparameter (the default setting in the GIFT toolbox is *a* = 0.5).

In our analysis, fifty three reproducible spatial references, denoted as *R*, were identified by analyzing two distinct fMRI datasets, namely the Genomics Superstruct Project (GSP) and the Human Connectome Project (HCP). The GSP dataset comprises 1005 subjects while the HCP dataset has 823 subjects [49]. These spatial references were derived using group ICA and are collectively referred to as the Neuromark fMRI 1.0 template. scICA is employed on the data matrix *X* by utilizing a fast fixed-point algorithm [49] which is incorporated into group ICA toolbox GIFT. One can think of this as applying a set of corresponding overlapping networks-of-interest to both PET and fMRI, where the spatially constrained ICA adapts the spatial priors to the data, resulting in PET specific and fMRI specific maps. We applied scICA on the PET and fMRI data separately. Prior work has shown that the scICA estimation approach does not overconstrain the solution and well captures individual variation.

### 2.3. Experiments with Spatially Constrained ICA

In this particular study, we analyzed data collected from 43 individuals who consented to participate in a study conducted at Emory University. Data from baseline and 1-year sessions were combined to maximize the amount of data used to create our dataset *X*. We first preprocessed the PET and fMRI data separately using the afore-mentioned preprocessing pipelines (see section 2.1). We then utilized scICA to analyze each preprocessed dataset separately. We provide a reference brain map *R* (see section 2.2) which contains the prior information of desired brain sources.

For PET data, the obtained results from scICA were decomposed into mixing and source matrices. The mixing matrix contains scores that show degree to which a given component is expressed in the data for a given subject. Specifically, the scores in the *i^th^*column represent each subject’s contribution to that component. On the other hand, the source matrix includes scores that identify distinct spatial patterns that indicate regions of coherent variability between subjects. The scores in each row of the source matrix correspond to different spatial configurations with statistical significance. For fMRI data, we obtained 53 group components from scICA. By applying back reconstruction techniques, we extracted subject-specific time-courses (or loading parameters in the case of PET) and sources. The overall procedure is shown in Figure 1.

**Figure 1:**
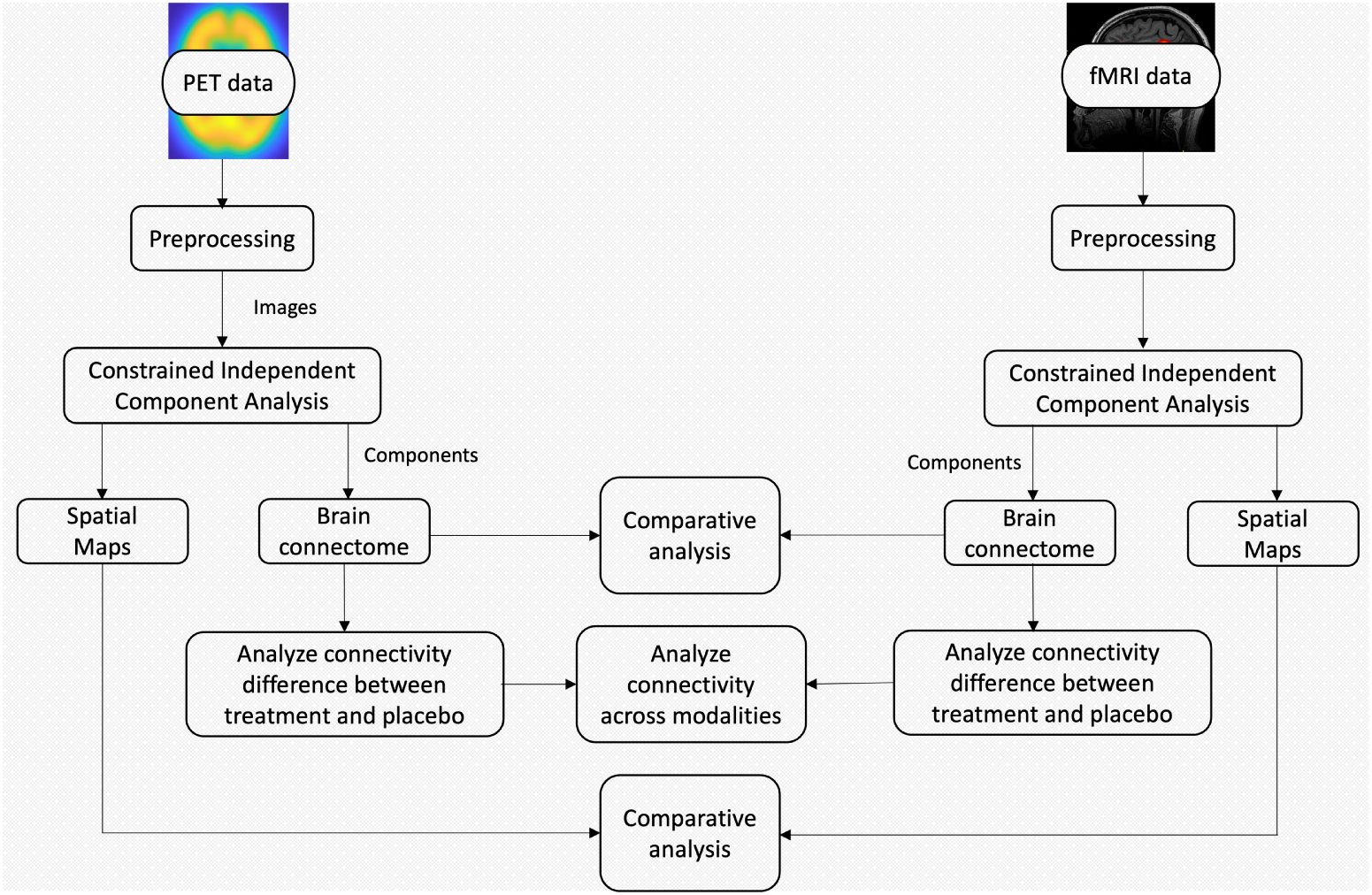
An overview of the proposed Neuromark PET approach. First, constrained ICA (cICA) was applied to the preprocessed PET and fMRI data separately. A comparative analysis was conducted between the brain connectomes and spatial maps (obtained from cICA) from the PET and fMRI modalities. Finally, connectivity differences were analyzed between the treatment and placebo groups.

### 2.4. Pairwise Correlations

We analyzed the obtained results from PET and fMRI data separately. For the PET data, we used the mixing matrix to calculate the pairwise correlations between the loading parameters. To better visualize the underlying molecular networks of the brain, 53 components resulting from this analysis were grouped into seven component regions : subcortical (SC), auditory (AUD), visual (VIS), sensorimotor (SM), cognitive control (CC), default-mode network (DMN), and cerebellar (CB). We note that these matrices reflect the covariation of the component among subjects. We consider this a type of inter-subject molecular network connectivity (covariation/coupling) in contrast to the intra-subject functional network connectivity as in the fMRI data.

For the fMRI data, we applied scICA to identify 53 group components. We then computed subject-specific time courses and spatial maps and evaluated functional network connectivity by calculating the pairwise correlations between each column of time courses. By taking the average of the functional network connectivity across all subjects, the mean FNC was computed. The 53 components were also partitioned into seven regions, following a similar process to the PET data analysis.

### 2.5. Group Connectivity Difference

In this study, we have MCI datasets divided into two groups: treatment and placebo, with 25 and 18 subjects, respectively. Our goal in this analysis was to evaluate the degree to which PET and fMRI might distinctly capture group differences. We did not investigate the goals of the clinical trial in this study. For the PET data, we employed the mixing matrix from the scICA outcomes to separate the loading parameters into two distinct groups. We subsequently calculated group-specific pairwise correlations and performed a two-sample t-test to determine the activation differences between the two groups. The activation difference was thresholded at *p<*0.05 after applying false discovery rate (FDR) correction.

For the fMRI data analysis, we similarly partitioned them into two groups: treatment and placebo. Initially, we calculated the functional network connectivity for each subject by computing the Pearson correlation between their respective time courses. Subsequently, we computed the average FNC for each group. To determine the differences between the two groups, we performed a two-sample t-test to observe the connectivity (FNC) differences between two groups. We then applied FDR correction at *p<*0.05 to identify statistically significant FNC differences.

## 3. Results

We present the whole brain connectome (MNC) of 53 components from the PET data in Figure 2(A). Each cell in the figure shows the correlation between loading columns *i* and *j*. The results indicate high positive correlations between most of the components of CON and DMN domains, suggesting strong positive connectivity (higher interaction between CON and DMN) between them. Conversely, the CBN domain demonstrated high negative correlations scores with both CON and DMN domains, indicating strong negative connectivity (anti correlation) between CBN and these domains. Furthermore, positive correlations were found between the CON and SMN domains, while the components within each domain exhibited highly positive correlations among themselves. The spatial maps of all 53 components are shown in Figure 2(B).

**Figure 2:**
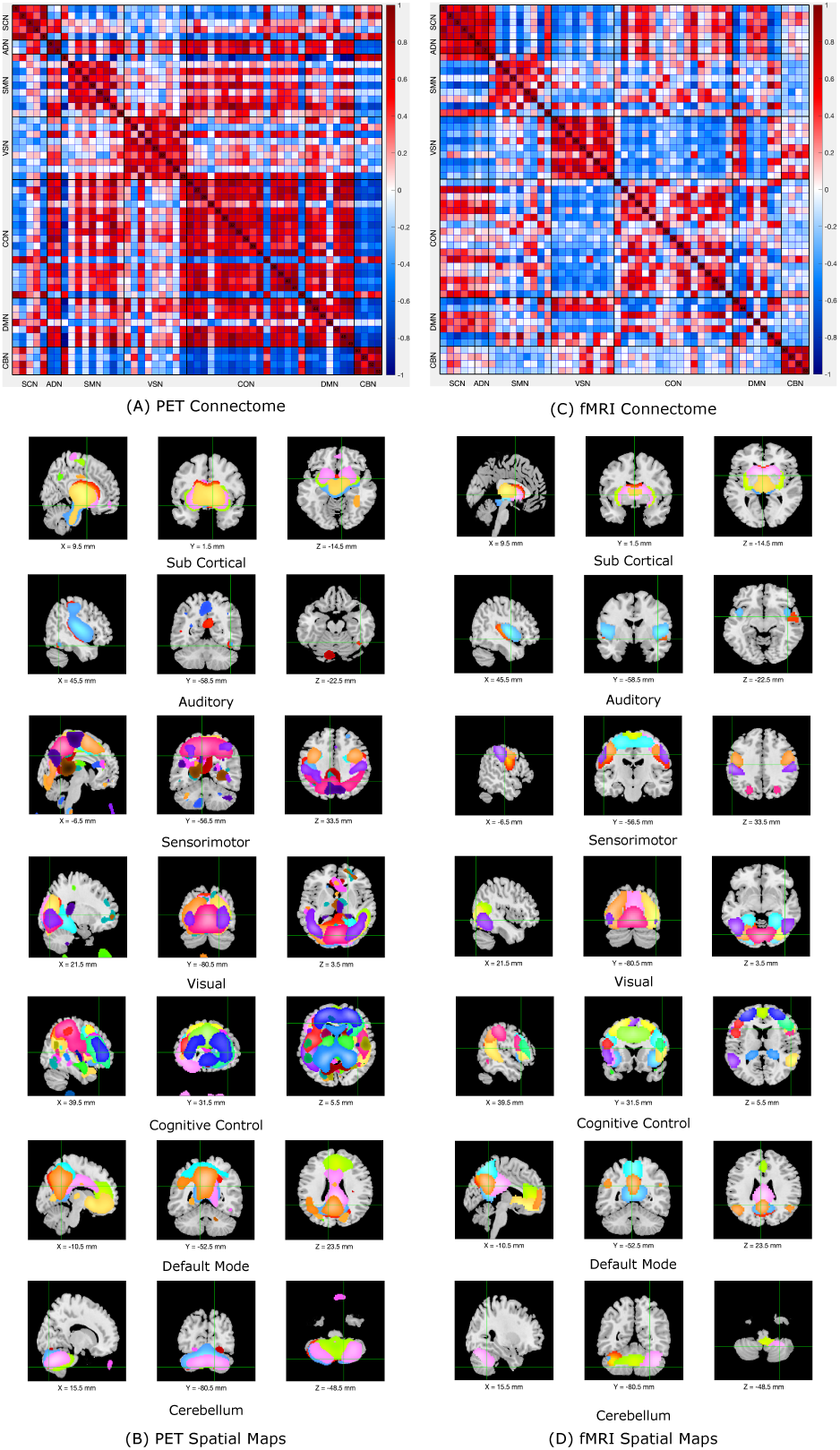
Whole brain connectome and spatial maps from PET and fMRI data. (A) and (B) represent the correlation between the loading parameters and the spatial maps of 53 components respectively from PET analysis. The positive and negative correlations are colored by the red and blue color respectively. (C) and (D) depict the functional network connectivity and spatial maps of 53 components respectively from fMRI analysis. Fifty-three components are divided into seven domains: subcortical, auditory, visual, sensorimotor, cognitive control, default-mode network, and cerebellar.

We also extracted 53 fMRI-specific components from the fMRI data using the spa-tially constrained ICA approach. Figure 2(C) shows the brain connectome derived from the fMRI data, with each cell representing the mean connectivity values between components *i* and *j*. We evaluated the subject-wise FNC and calculated the mean FNC across all subjects. The results revealed higher positive correlation scores between the ADN and SCN domains, indicating strong positive connectivity (higher interactions) between them. Positive functional connectivity was also observed between the CON and SCN domains, and some of the components exhibited positive functional network connectivity between the DMN and SCN domains. The VSN domain demonstrated a high negative correlation score with the SCN domain, indicating strong negative connectivity (anti correlations) between them. Additionally, negative functional network connectivity was observed between the CON and VSN domains. Finally, the components within each domain (except for DMN) exhibited high positive connectivity among themselves. The spatial maps for all the components are presented in 2(D).

The analysis in Figure 2 reveals significant modular similarity between the PET and fMRI connectomes, with some differences specific to each modality. Specifically, the PET connectome shows strong positive connectivity between the CON to DMN and CON to SMN, while the fMRI connectome shows weak negative connectivity across these domains. In contrast, the CBN to CON and CBN to DMN exhibit strong negative connectivity in the PET connectome, whereas the fMRI connectome displays only weak negative connectivity. Furthermore, Figure 2(B) illustrates strong positive functional connectivity between ADN to SCN in the fMRI connectome but not in the PET connectome. Lastly, negative functional connectivity between CON to VSN is evident in the fMRI connectome but not observed in the PET connectome.

We also analyzed the spatial similarity between the PET and fMRI spatial maps. Figure 3(A) and (B) represent the within modality spatial correlation among the 53 components of either the PET and fMRI source matrices respectively. Figure 3(C) represents the inter-modality cross-correlations between the components from PET and fMRI networks. The correlations across the diagonal represent how similar or dissimilar PET and fMRI components exhibit. These diagonal correlations are presented in a histogram in Figure 3(D). We identify eight most correlated components (Correlation range : 0.65 *≤ corr ≤* 0.76). We also extracted eight least correlated components (correlation range : 0.17 *≤ corr ≤* 0.39). We present eight most and least correlated components in Figure 4(A) and (B) respectively.

**Figure 3:**
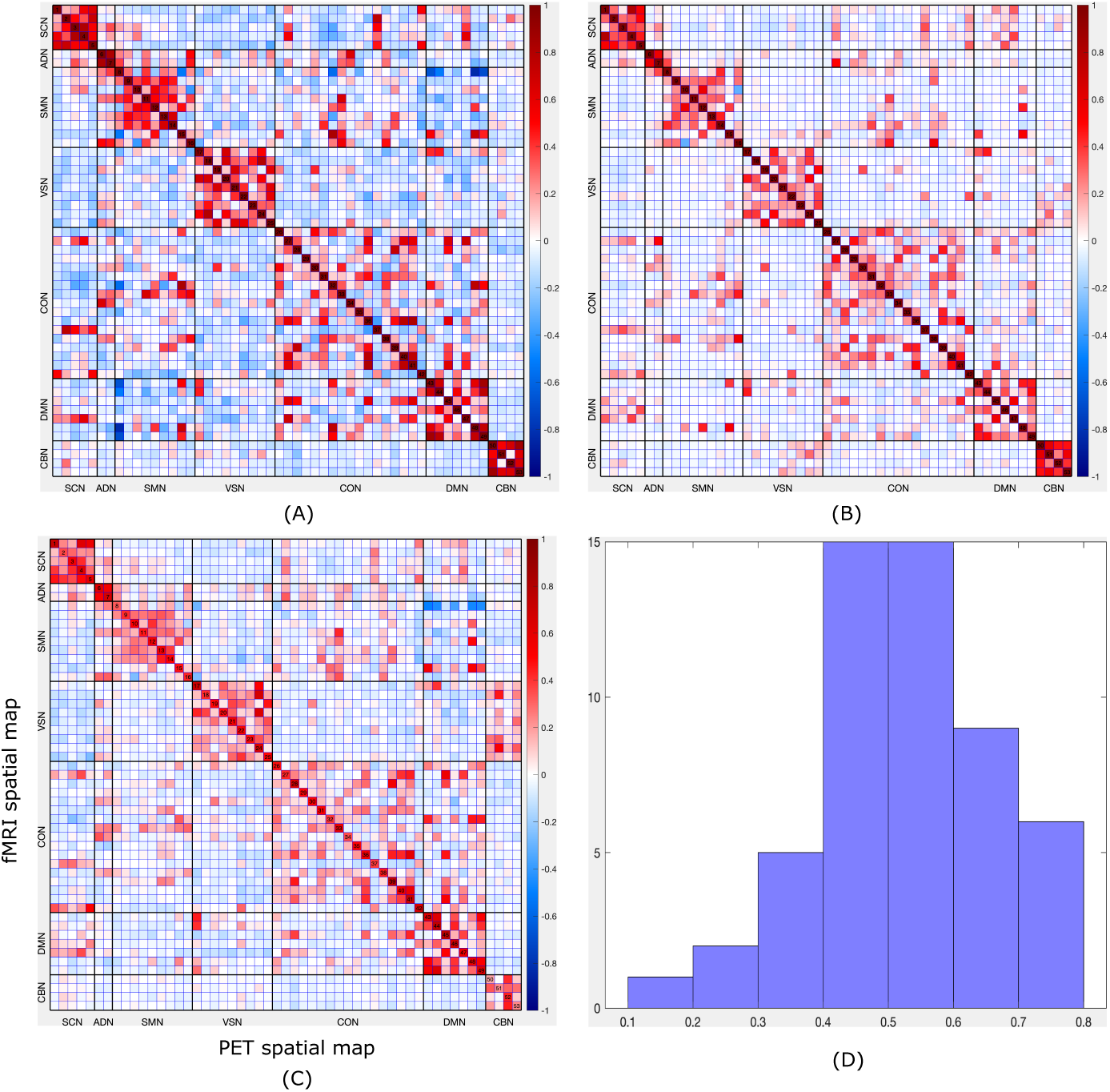
PET and fMRI spatial map correlations. (A) and (B) represent the within modality correlation among the spatial maps of 53 components from PET and fMRI analysis respectively. (C) represents the between-modal spatial correlations between the PET and fMRI spatial maps and these correlations are presented in histogram plot (D) to highlight the variation in spatial maps between the two modaliites.

**Figure 4:**
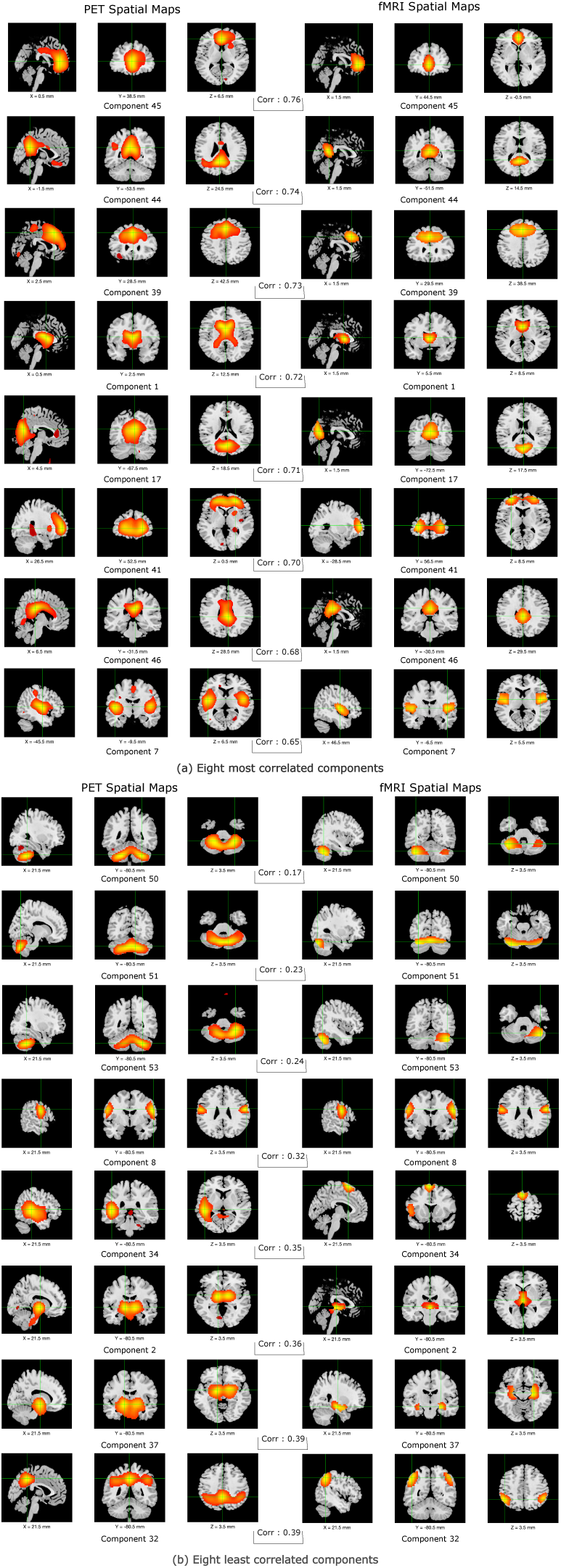
Most and least correlated components from PET and fMRI analysis.

The results of our study comparing the connectivity differences in individuals with MCI who received either medication or a placebo are presented in Figure 5. In this figure, a positive score indicates that individuals receiving medication had higher connectivity differences compared to those in the placebo group, while negative scores indicate components in the placebo group that had higher connectivity. The connectivity differences from PET and fMRI analyses are shown in Figures 5 (A) and (B), respectively. The PET analysis showed that some components in those receiving medication had a higher connectivity difference compared to the placebo group in the SMN-SCN, CON-SCN, CON-SMN, and DMN-SCN domains. In contrast, the placebo group had higher connectivity difference compared to medication in the VSN-SMN and CBN-VSN domains. Meanwhile, the fMRI analysis showed higher connectivity differences for those receiving medication compared to placebo in the VSN-SCN and CON-VSN domains, while placebo subjects had higher connectivity differences in the ADN-SCN and CON-SMN domains. We also calculated the common connectivity difference from both PET and fMRI analyses and presented the results in Figure 5(C). We found a single component in the CON-SCN, CON-SMN, DMN-SMN domains, and two components in the SMN domain that showed common connectivity differences in both modalities.

**Figure 5:**
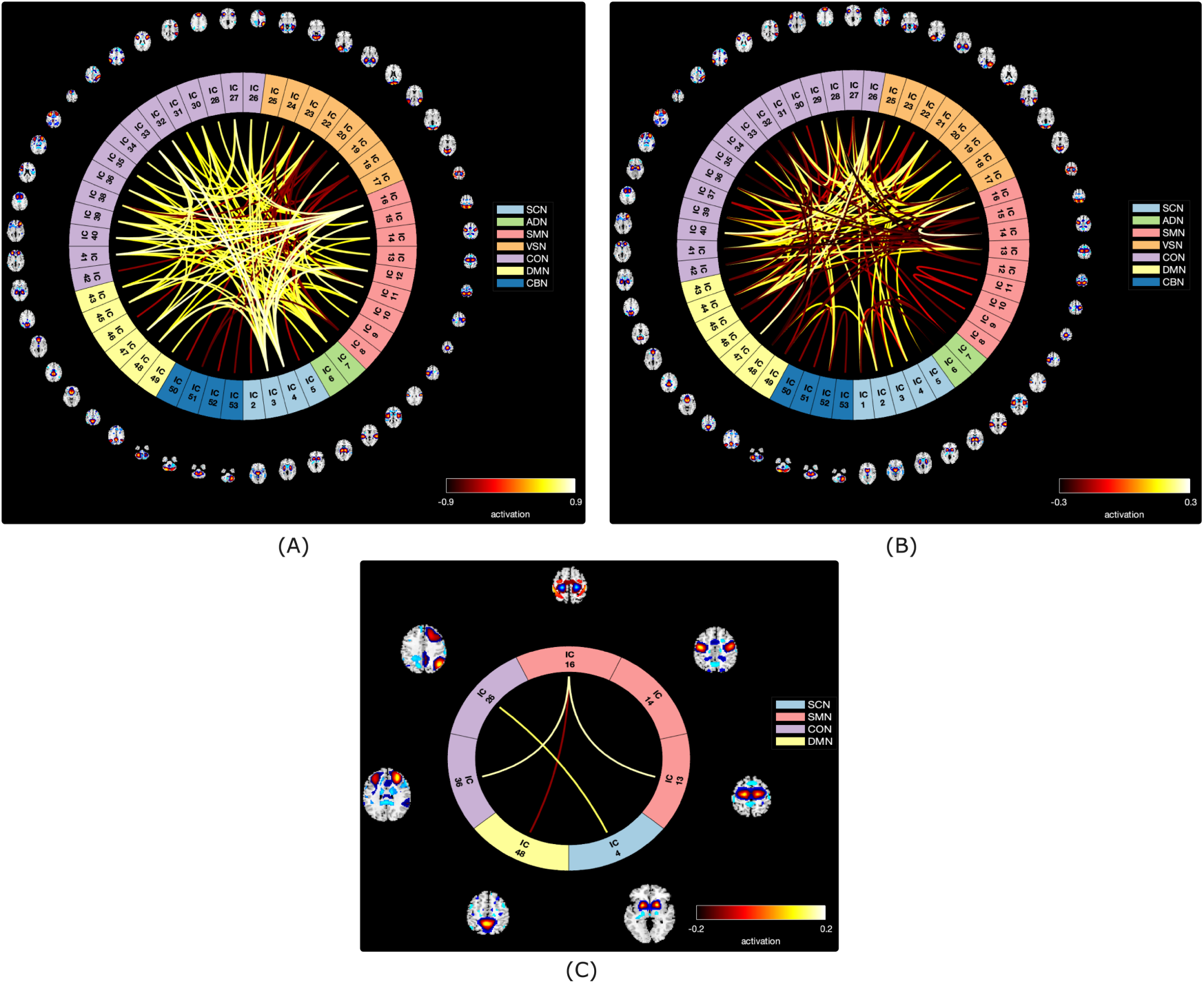
Connectogram maps of PET and fMRI brain connectivity differences. (A) and (B) represent the brain connectivity difference between the subjects who took candesartan and the subjects with placebo for the PET and fMRI analysis respectively. (C) represents common connectivity difference between treatment and placebo from PET and fMRI modalities and this result is generated after the element-wise multiplication between (A) and (B)

## 4. Discussion

In this study, we utilized a scICA technique to analyze data obtained from both PET and fMRI imaging modalities. Our results indicate the presence of modularized PET connectomes that complement those identified using resting-state fMRI. Additionally, we observed distinguishable brain networks identified from the result estimates obtained from PET and fMRI. We utilized the same NeuroMark 53 network templates described in [50] for our analysis and present in Figure 6.

**Figure 6:**
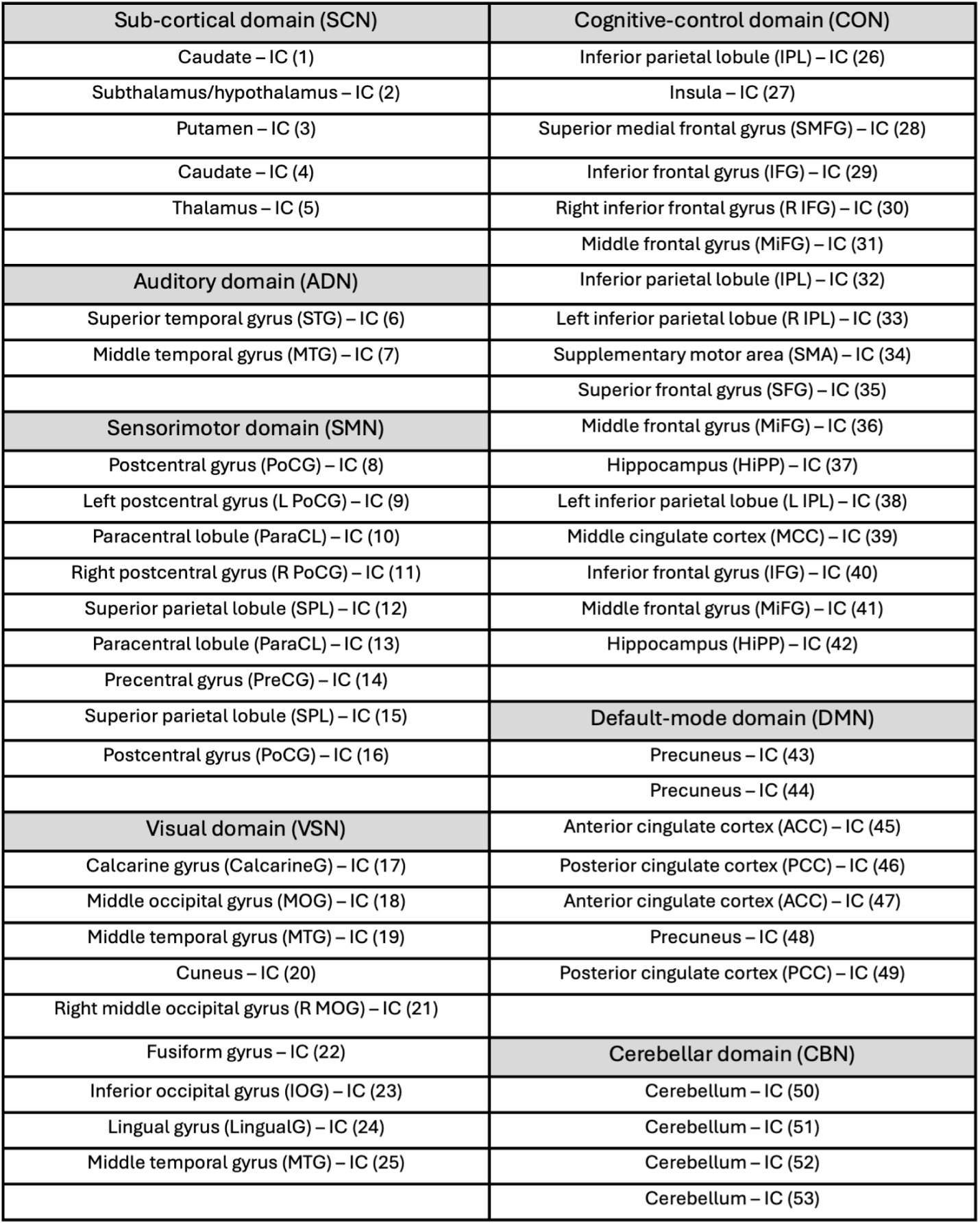
Information of the extracted network templates. Here, each network template is represented by one independent component (IC)

In fMRI, functional network connectivity (obtained from fMRI time series) is commonly utilized to measure the strength of connections among various brain networks. Recent studies have examined changes in brain functional (network) connectivity among patients with MCI and AD, using whole brain or region of interest approaches [17, 51, 52]. The subcortical region, such as the thalamus, is of particular interest in such studies given its widespread physical connection with most cortical areas. In a study by Wang et al., rsfMRI was used to investigate changes in thalamic connectivity in patients with MCI [53]. The study revealed regions that exhibited decreased functional connectivity with the thalamus, including the fusiform gyrus (FG), middle occipital gyrus, and bilateral cuneus in the visual domain; the superior frontal gyrus (SFG) in the cognitive control domain; the paracentral lobe (PCL), superior parietal lobe (SPL), and precentral gyrus in the sensorimotor domain; and the precuneus in the default mode domain. Our own fMRI and PET studies similarly identified negative functional and molecular connectivity respectively between the thalamus and these regions. These findings suggest that both fMRI and PET modalities capture changes in thalamic connectivity, albeit linked to different brain regions.

Other research has demonstrated a link between disrupted intrinsic connectivity and episodic memory (EM) deficits in individuals with amnestic mild cognitive impairment (aMCI) [54]. The functional integration of insula networks has been identified as an important factor in the EM process. Previous studies have shown that insula atrophy is a reliable marker of AD development, distinguishing AD patients from healthy individuals [55]. Studies by Xie et al. have found the disrupted intrinsic connectivity of insula networks (in cognitive control domain) with the frontal, parietal, and temporal occipital lobes(in visual domain), and subcortical regions. Our fMRI analysis revealed strong anticorrelations between the insula and middle occipital gyrus, Right middle occipital gyrus and Inferior occipital gyrus. From our PET analysis, we observed strong anticorrelations between insula and Middle occipital gyrus and weak correlations with right middle occipital gyrus and inferior occipital gyrus. We also observed anticorrelations between the VIS and AUD domains. Similar findings are reported in [54] where decreased anticorrelated connectivity were found in the VIS and AUD pathway in aMCI patients.

Abnormal connectivity between the insula and other brain networks may alter the semantic memory network, as described in recent research, and could result from compensatory mechanisms [56]. Our fMRI analysis identified more disruptive connectivity between the insula and other brain networks compared to PET studies. Given the association between insula networks and high-order cognitive processes, further investigation of the disruptive connectivity with the insula in MCI individuals should be pursued in future studies.

In the past two decades, numerous studies have indicated that alterations in the functional and structural connectivity within the DMN serve as significant indicators of AD [57, 58, 59]. Our analysis revealed strong intra-domain connectivity in the DMN based on PET imaging. Vincent et al. investigated changes in connectivity over time in individuals with MCI who either progressed to dementia or remained clinically stable [60]. They observed higher DMN connectivity in the dementia group compared to the stable MCI group during the final MRI session. These findings provide evidence that the strong intra-domain connectivity in the DMN is not a random occurrence. Surprisingly, from PET study, We also observed strong connectivity between the DMN and cognitive network domain. Recent studies has observed that there was considerable spatial overlap of *Aβ* accumulating regions with cognitive control domain in AD/MCI patients [61]. We speculate that there lies some underline connectivity between DMN and CON domain which is strongly visible in the PET analysis. Notably, we did not observe strong connectivity like the above in the fMRI analysis suggesting enhanced PET sensitivity relative to fMRI for some connections.

The latest research has established that the cerebellum, traditionally associated with motor control, is now recognized as a center for higher cognitive and emotional functions, even in the developing brain [62, 63]. Various studies are currently being conducted to investigate the connectivity between the cerebellum and the cognitive control domain in order to identify biomarkers in patients with MCI [64, 65]. Tang et al. discovered a decrease in connectivity between the CBN and CCN, such as the superior frontal gyrus and inferior frontal gyrus, in MCI patients. Bai et al. also reported the negative connectivity between the CBN and frontal gyrus [66]. Our PET analysis (Figure 2(A)) also revealed strong negative connectivity between the CBN and CCN domains, while showing increased positive connectivity between the CBN and the hippocampus. Similar findings of increased connectivity between the hippocampus and CBN have been reported in ischemic stroke patients, who are at a higher risk of developing vascular cognitive impairment [67]. Additionally, our analysis also identified strong negative connectivity between the CBN and DMN. Furthermore, fMRI analysis has also shown negative connectivity between CBN and CON as well as DMN, although this connectivity is more pronounced in the PET analysis.

We also identified significant connectivity differences between MCI patients who received either medication or a placebo in both modalities. The fMRI analysis (Figure 5(B)) revealed higher connectivity in the medication group between the thalamus (from SCN) and the middle occipital gyrus (from VSN). However, this connectivity was not observed in our PET analysis. In the fMRI group fMRI study, we found strong negative connectivity between the insula and the right middle occipital gyrus (Figure 2(B)). Interestingly, MCI patients with medication showed higher connectivity between these components (Figure 5(B)). Additionally, in the group analysis from both PET and fMRI studies, we observed strong negative connectivity between the CBN and CON. However, in our fMRI analysis, we found higher connectivity between the CBN and the left inferior parietal lobe, as well as the supplementary motor area, in individuals treated with medication. In PET analysis, we found higher connectivity between the anterior cingulate cortex and the left inferior parietal lobe; and for fMRI studies we got higher connectivity between the anterior cingulate cortex and the insula, precuneus and superior medial frontal gyrus, precuneus and middle frontal gyrus, precuneus and middle cingulate cortex, and precuneus and inferior frontal gyrus in individuals treated with medication. These findings suggest that treated MCI patients exhibited increased connectivity in the aforementioned brain regions/components, despite showing negative connectivity prior to treatment. Furthermore, both PET and fMRI analyses revealed common connectivity between the postcentral gyrus and paracentral lobule, postcentral gyrus and precentral gyrus, inferior parietal lobule and caudate, middle frontal gyrus and postcentral gyrus, and precuneus and postcentral gyrus. However, we observed higher connectivity in the CON and SMN domains in PET analysis in MCI patients treated with medication. Conversely, we noticed mild activation in MCI patients treated with a placebo in the fMRI study. Overall PET and fMRI in this case appear to be more complementary than common in the group results, further supporting the importance of using a multimodal approach. The main goal of this analysis was to examine similarities and differences in the rest fMRI and PiB PET changes. To evaluate this we tested for differences in patients taking medication compared to those on a placebo. We did not focus on the specific goals of the clinical trial in this analysis.

## 5. Limitations and Future Work

In our analysis, we used rest fMRI networks estimated from independent data to spatially constrain, but still adapt to, the PET data, to estimate covarying PET networks corresponding to the rest fMRI network. This provide a common inferential framework and allows us to evaluate the structure of the connectome between modalities. There are several reasons for this, resting networks in fMRI are widely studied and replicated in multiple studies, and secondly there is plenty of resting fMRI data to generate independent templates. However we can also take a similar approach to PET data. In future work, we plan to bring in additional data to generate a PET-specific template and used this to constrain the fMRI data and compare the approaches. Throughout the study, we employed ICA under the assumption of linearity to extract resting-state networks. Nevertheless, a recent study conducted by Motlaghian et al. highlighted the presence of nonlinear relationships within functional networks, a factor often disregarded in linear analyses [68]. Exploring nonlinear methodologies could prove valuable in examining the comprehensive brain connectomes among individuals with MCI. Due to the relatively small sample size, we used all data for the study to ensure a robust decomposition, though all statistical comparisons are performed on independent subjects. Future studies should work with larger sample sized to replicate and extend our results. In this study we also estimated group differences between medicated versus placebo groups at baseline. This was done to evaluate the feasibility of the proposed framework to capture group differences, and was not designed to be a clinical trial. The approach however is very general and we suggest may be a powerful tool for studying PET/fMRI relationships in a wide variety of future studies.

## 6. Conclusion

PET and fMRI are two commonly used neuroimaging methods for investigating brain activity. Recent studies have utilized both PET and fMRI connectomes to analyze brain connectivity. However, most studies do not use data-driven approaches to estimate networks and compare whole brain connectomes in a high dimensional network space. To address these limitations, this research paper introduces a novel data-driven constrained ICA approach with a prior reference brain template. To our knowledge, this study presents the first comparison of whole brain connectomes using both PET and fMRI imaging modalities. Results demonstrate significant modular similarity between the PET and fMRI connectomes, suggesting that both methods are capturing certain aspects of the modularity of brain activity. Moreover, modality-specific connectivity was also discovered. Furthermore, we also analyzed the differences among participants with MCI who received either medication or a placebo. The findings revealed both common and modality specific treatment effects on the brain connectomes, as identified by fMRI and PET imaging modalities. The study concludes that combining PET-specific resting-state network with fMRI provides a means of identifying complementary information in both modalities and creates new opportunities for developing biomarkers.

## Acknowledgements

This study was funded by NSF (2112455) and NIH (R01AG073949).

## Footnotes

1 http://afni.nimh.nih.gov/

## Notes

### Competing Interest Statement

The authors have declared no competing interest.

### Summary of Updates

This version of the manuscript has been revised to update Figures 1 and 5, and the paper has been edited primarily in the Introduction, Results, and Discussion sections."

